# Structural Basis for the Calmodulin-Mediated Activation of eEF-2K

**DOI:** 10.1101/2022.01.15.476372

**Authors:** Andrea Piserchio, Eta A. Isiorho, Kimberly Long, Amanda L. Bohanon, Eric A. Kumar, Nathan Will, David Jeruzalmi, Kevin N. Dalby, Ranajeet Ghose

**Author notes:** Current Address: Laboratory of Molecular Electron Microscopy, the Rockefeller University, New York, NY 10065.

## Abstract

Translation is a highly energy consumptive process^1^ tightly regulated for optimal protein quality^2^ and adaptation to energy and nutrient availability. A key facilitator of this process is the α-kinase eEF-2K that specifically phosphorylates the GTP-dependent translocase eEF-2, thereby reducing its affinity for the ribosome and suppressing the elongation phase of protein synthesis^3,4^. eEF-2K activation requires calmodulin binding and auto-phosphorylation at the primary stimulatory site, T348. Biochemical studies have predicted that calmodulin activates eEF-2K through a unique allosteric process^5^ mechanistically distinct from other calmodulin-dependent kinases^6^. Here we resolve the atomic details of this mechanism through a 2.3 Å crystal structure of the heterodimeric complex of calmodulin with the functional core of eEF-2K (eEF-2K_TR_). This structure, which represents the activated T348-phosphorylated state of eEF-2K_TR_, highlights how through an intimate association with the calmodulin C-lobe, the kinase creates a “spine” that extends from its N-terminal calmodulin-targeting motif through a conserved regulatory element to its active site. Modification of key spine residues has deleterious functional consequences.

## Introduction

Eukaryotic protein levels are predominantly controlled by mRNA translation^7^, an energetically expensive process^1^ that is highly regulated^8^ to ensure optimal transit times and maintain protein quality^2^, and fashion the cellular response to environmental stress and changes in energy/nutrient availability^9^. A primary driver of this regulation is the specific phosphorylation (on Thr-56) of the GTP-dependent translocase, eukaryotic elongation factor 2 (eEF-2). This covalent modification, uniquely catalyzed by the α-kinase^10^, eukaryotic elongation factor 2 kinase (eEF-2K)^11,12^, diminishes the ability of eEF-2 to engage the ribosome suppressing the elongation phase of protein synthesis^3,4^. Dysregulation of eEF-2K activity has been linked to neurological conditions such as Alzheimer’s-related dementia^13^ and Parkinson’s disease^14^. Aberrant eEF-2K function has been correlated to enhanced tumorigenesis^15^, invasion, and metastasis^16^. eEF-2K is, therefore, an emerging target for the development of therapeutics against several diseases, including many cancers^17^, and neuropathies^18^, underscoring the importance of understanding its regulation in mechanistic detail.

eEF-2K is the only α-kinase reliant on calmodulin (CaM)^11^ for activation through an allosteric process^5^. CaM binding enhances the apparent *k*_cat_ towards a peptide-substrate by ~2400-fold without significantly altering affinity. Subsequent auto-phosphorylation at the primary upregulating site (T348) provides an additional ~6-fold increase in *k*_cat_, yielding the fully activated state. This mechanism is distinct from other CaM-dependent kinases that rely on CaM-induced displacement of an auto-inhibitory segment for enhanced substrate access^6^. eEF-2K activity is further modulated by Ca^2+19^, pH^20^, and regulatory phosphorylation^21^ via multiple pathways, including mTOR, AMPK, and ERK^15^. Obtaining insight into eEF-2K activation and regulation in atomic detail has been hindered by the absence of structures of the full-length enzyme or its CaM-bound activated complex. We have identified a minimal functional eEF-2K construct, eEF-2K_TR_ (Fig. 1a, Extended Data Fig. 1), activated by CaM similarly to the wild-type enzyme and efficiently phosphorylates eEF-2 in cells^22^. Using X-ray crystallography, we resolved the structure of its T348-phosphorylated state (*p*eEF-2K_TR_) in an activated heterodimeric complex with CaM (CaM•*p*eEF-2K_TR_) to 2.3 Å. Almost four decades after its discovery and characterization^11^, this structure provides the first critical insight into the atomic details of the unique mechanism of the CaM-mediated activation of eEF-2K.

**Fig. 1.**
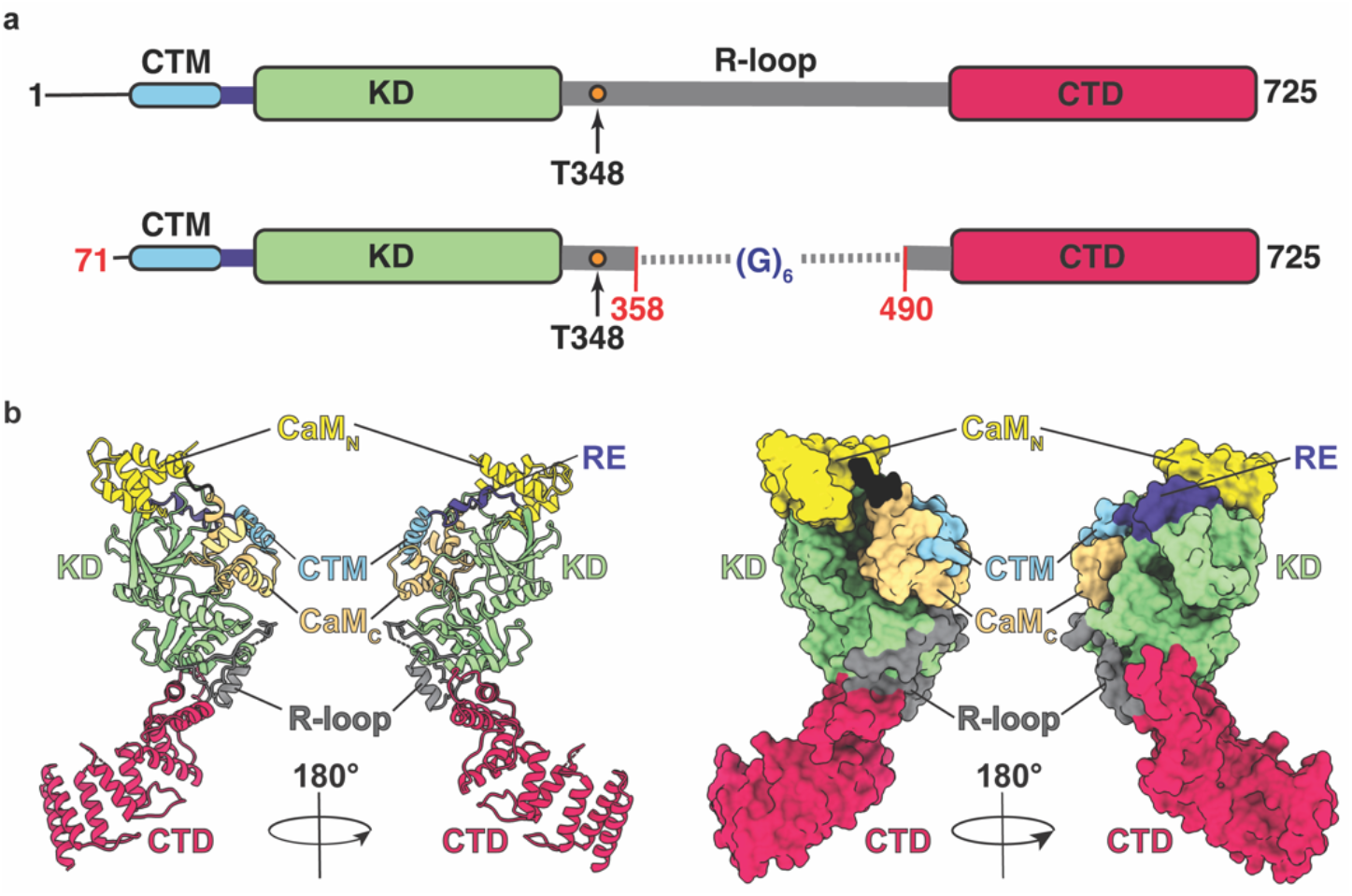
Structure of the CaM•*p*eEF-2K_TR_ complex. **a**, Schematic representation of the eEF-2K (top) structural modules indicating the CTM (cyan), RE (navy-blue), KD (lime-green), R-loop (grey) and CTD (crimson). eEF-2K_TR_ (bottom) is missing 70 N-terminal residues, and a 6-glycine linker replaces the 359-489 segment of the R-loop. **b**, Structure of the CaM•*p*eEF-2K_TR_ complex shown in ribbon (left) or surface (right) representation; CaM_N_ and CaM_C_ are colored bright and dull yellow, respectively. The same color scheme is used in all figures unless otherwise stated.

## Results and discussion

### Structural features of *p*eEF-2K_TR_

The CaM•*p*eEF-2K_TR_ complex forms an elongated structure (Fig. 1b) consistent with previous solution-state X-ray scattering (SAXS) measurements on the inactive complex (unphosphorylated T348)^22^ (Extended Data Fig. 2). The kinase domain (KD) shows a characteristic α-kinase architecture^23,24^ with the “catalytic” residues in orientations suggestive of an active conformation (Extended Data Fig. 3a). While the P- and N/D-loops are closed (Extended Data Fig. 3b), as in the nucleotide-bound states of the related MHCK-A KD^24,25^, no density corresponding to nucleotide was apparent despite crystallization with the ATP-analog, AMP-PNP. Phosphorylated T348 (*p*T348; Extended Data Fig. 4) docks into a phosphate-binding pocket (PBP) hydrogen-bonding with the K205, R252, and T254 side-chains, and the L253 and T254 main-chains (Fig. 2a). D280 on the catalytic-loop hydrogen bonds with the side-chains of R252 and Y282. This interaction promotes methyl-*π* interactions between I275 and Y282 (Fig. 2a), thereby coupling the catalytic site to the PBP.

**Fig. 2.**
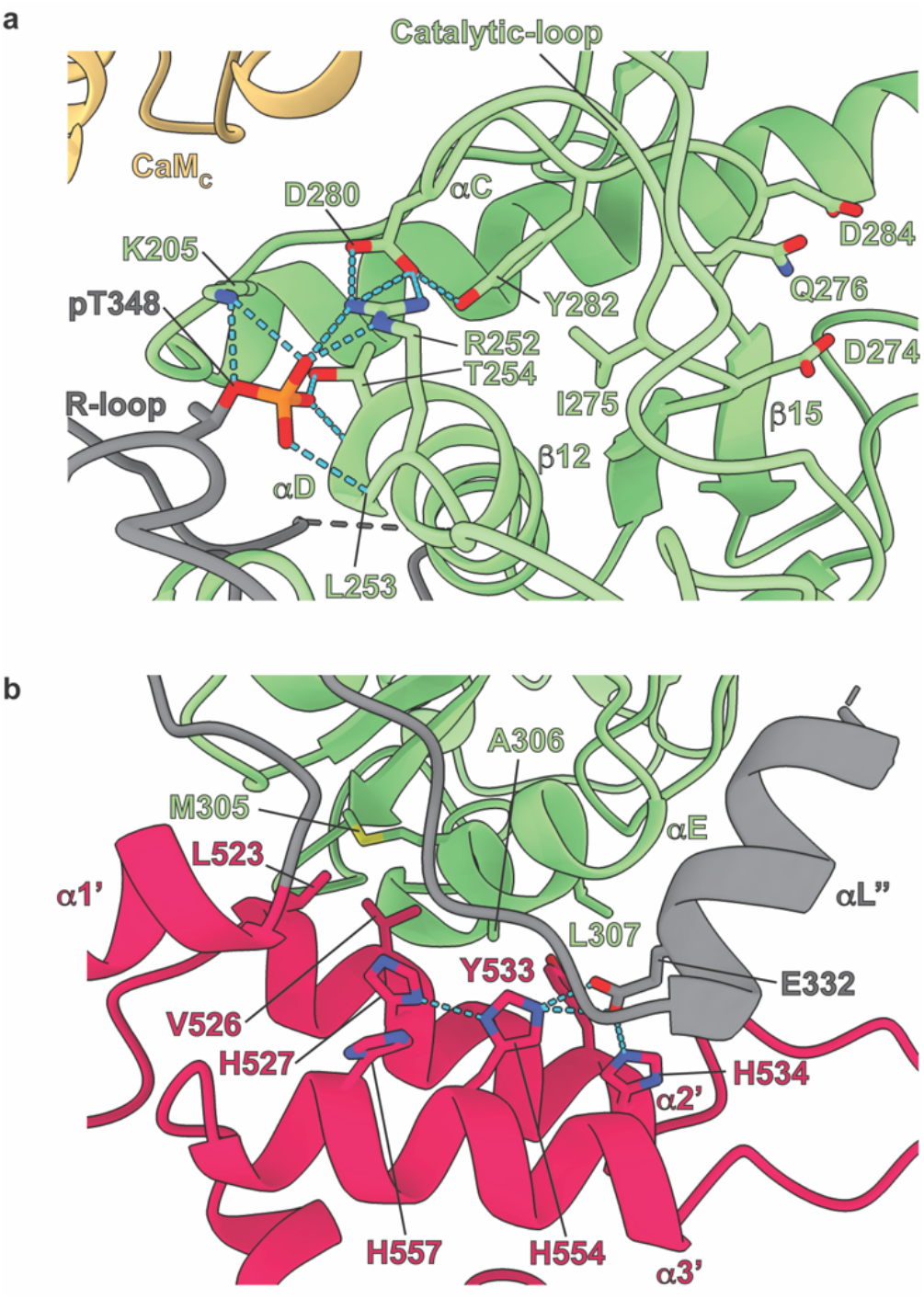
Intramolecular interactions within *p*eEF-2K_TR_ in the CaM•*p*eEF-2K_TR_ complex. **a**, Interactions that stabilize *p*T348 at the phosphate-binding pocket (PBP) and couple the PBP to the active site are indicated. Key catalytic-loop residues, including the base D274, Q276, and D284, are shown for reference. **b**, Key interactions involving the αE-α2’-α3’ element that stabilize the KD/CTD interface are shown.

The all-helical C-terminal domain (CTD) represents a feature unique to eEF-2K that forms a docking platform for eEF-2^26,27^. The α2’-α7’ region displays a pattern resembling SEL1-like repeats^28^, the α8’-α10’ segment deviates from this arrangement, and α1’ is somewhat disengaged from the rest of the CTD. α2’ stabilizes the KD/CTD interface, interacting with αE on the KD C-lobe (KD_C_) and α3’ (Fig. 2b). The αE-α2’ interface features hydrophobic interactions involving Y533 sandwiched by A306 and L307, and V526 inserted between M305 and L523. A curious feature within the αE-α2’-α3’ element is the linear spatial positioning of three conserved His residues (H527 and H534 on α2’; H554 on α3’; Fig. 2b); the also spatially proximal H557 stacks with H527. H554 forms a rare δ-δ hydrogen bond with H527^29^. H534 and H554 form salt bridges with the conserved E332 on a helical R-loop segment (αL’’). It is tempting to speculate that alternative protonation states of one or more of these His residues would modify interactions within the KD/CTD interface, and perhaps its coupling to the R-loop, through the αE-α2’-α3’ element, and thereby contribute to the observed pH response of the enzyme^20^. NMR studies^27^ have localized the eEF-2 docking-site to the C-terminus of the CTD (α9’-α10’), ~60 Å away from the active site, suggesting that distinct interactions drive eEF-2 recognition and its subsequent phosphorylation.

### Recognition of *p*eEF-2K_TR_ by CaM

CaM contacts *p*eEF-2K_TR_ in an extended conformation using both its N-(CaM_N_) and C-terminal (CaM_C_) lobes, with CaM_C_ being more intimately associated with the kinase (Fig. 3a). This assembly is consistent with changes in protection seen in previous hydrogen exchange mass spectrometry (HXMS) analyses (Extended Data Fig. 5)^22,30^. All four CaM_C_ helices engage the CaM-targeting motif (CTM) through their hydrophobic faces in a Ca^2+^-bound “open” conformation, and all, except α_H_, interact with the N-lobe of the KD (KD_N_) using their predominantly hydrophilic faces. A deep hydrophobic pocket, lined by the side-chains of Ala88 (three-letter codes used for CaM residues), Phe92, Ile100, Leu105, Met124, Ile125, Val136, Phe141, Met144, and Met145, accommodates W85 from the eEF-2K_TR_ CTM. Additional hydrophobic interactions of I89 with Ala88 and Met145, and of A92 with Val91 and Leu112 generate a 1-5-8 binding mode (Fig. 3b) within a helical CTM (αCTM) like in the NMR structure of the complex of CaM with a peptide encoding the CTM (CTM-pep, 74-100)^31^. However, no CaM_C_-bound Ca^2+^ ions were seen in the peptide complex^31^. The hydrophilic face of CaM_C_ makes numerous contacts with KD_N_ over a large interaction surface (Fig. 3c). A “virtual alanine scan” at the interface identified several contacts involving residues for which *in silico* Ala mutations are substantially destabilizing (ΔΔG > 1.5 kcal/mol)^32^. These include Asp95 and Ser101 on CaM_C_ and K192 and D208 on the eEF-2K_TR_ KD, whose side-chains form a mesh of hydrogen bonds centered on K192 (ΔΔG=2.5 kcal/mol). Other notable interactions at the interface include a hydrogen bond between the Arg90 side-chain and the A164 main-chain and an L281-His107 methyl-*π* interaction. These interactions collectively localize the CTM-tethered CaM_C_ behind the kinase active site. An additional salt bridge between Asp122 and R351 couples CaM_C_ to the R-loop.

**Fig. 3.**
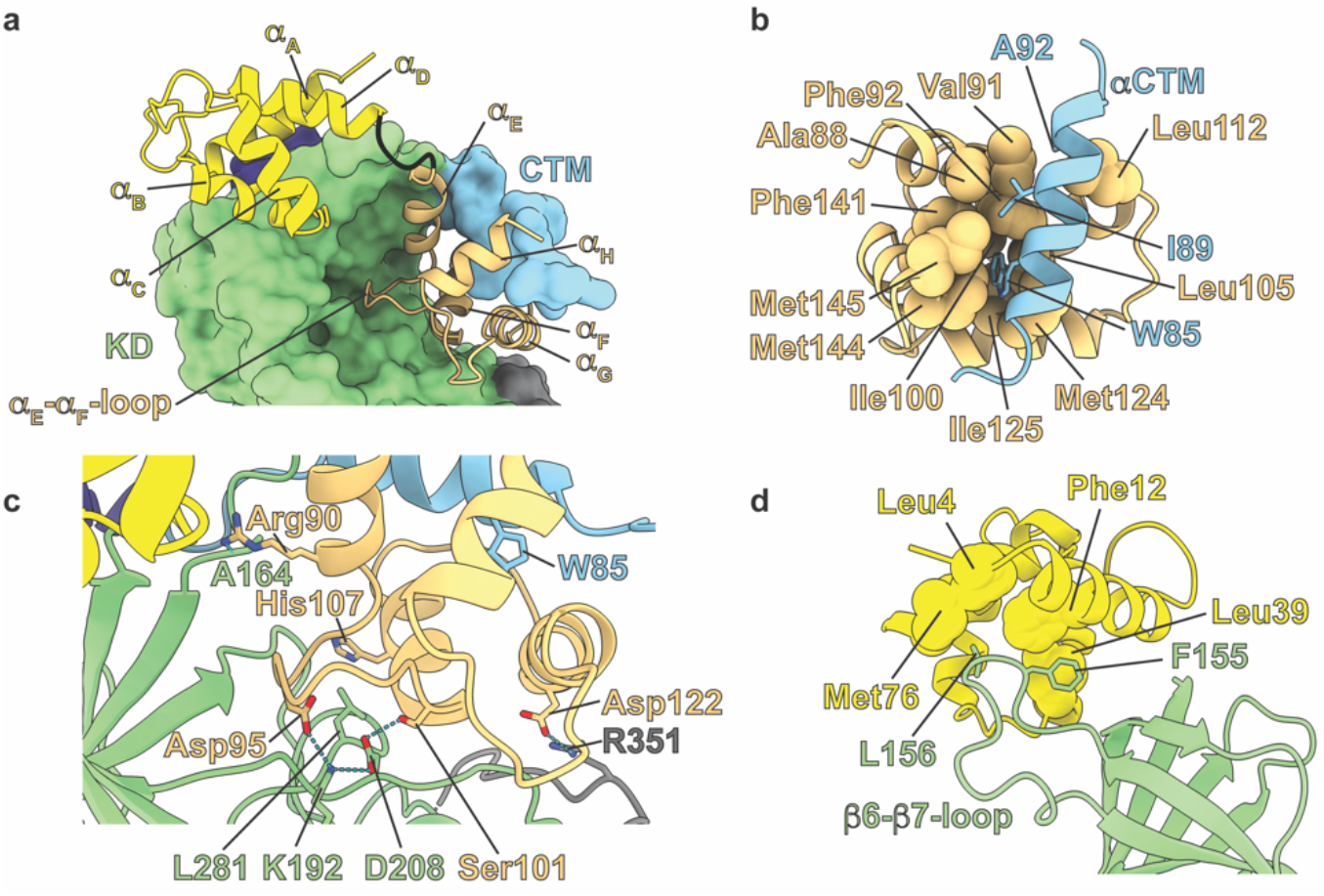
Intermolecular interactions between CaM and *p*eEF-2K_TR_. **a**, CaM and eEF-2K_TR_ are shown in ribbon and surface representations, respectively. CaM_N_ (α_A_-α_D_) and CaM_C_ α_E_-α_G_) helices are indicated. Interactions with the hydrophobic **(b)** and hydrophilic **(c)** faces of CaM_C_ with the CTM and KD_N_, respectively. **d**, Interactions of CaM_N_ with the β6-β7-loop of KD_N_. Elements not directly involved in the interaction are hidden to aid visualization.

CaM_N_ engages *p*eEF-2K_TR_ in a Mg^2+^-bound “closed” state^33^ contrasting its Ca^2+^-bound “open” conformation seen in the CTM-pep complex (Extended Data Fig. 6a,b)^31^. To the best of our knowledge, this conformation has not been previously observed in a CaM complex. In this configuration, the KD_N_ β6-β7-loop associates with shallow hydrophobic grooves on CaM_N_; F155 interacts with Leu39 and Phe12, and L156 with Leu4 and Met76 (Fig. 3d). CaM_N_ is further stabilized through numerous crystal contacts with symmetry-related neighbors (Extended Data Fig. 6c). Given this unique conformation and the high Mg^2+^concentration for crystallization, we tested the influence of Mg^2+^ on CaM_N_/*p*eEF-2K_TR_ interactions in solution. While the concentration of free Mg^2+^ in mammalian cells remains relatively constant at ~1 mM^34^, the Ca^2+^ concentration, ~50-100 nM in resting cells, can be enhanced to ~100 μM^35^ in calcium-microdomains under specific stimuli. The Ca^2+^:Mg^2+^ ratio (1:100) used for crystallization lies well within this cellular range. eEF-2K_TR_ retains the ability to efficiently auto-phosphorylate on T348 under these conditions (Extended Data Fig. 6d). NMR analyses (Extended Data Fig. 7) using Ile-δ1 and Met-*ε* resonances of CaM indicate no appreciable perturbations in the CaM_C_ resonances with increasing Mg^2+^ within a complex formed in the presence of Ca^2+^. This suggests that Mg^2+^, even at very high concentrations, is unable to affect the interactions of *p*eEF-2K_TR_ with CaM_C_ significantly. In contrast, CaM_N_ resonances display evidence of exchange between Ca^2+^- and Mg^2+^-bound conformations with increasing Mg^2+^. Indeed, at very high Mg^2+^ concentration where the Ca^2+^:Mg^2+^ ratio is heavily skewed towards Mg^2+^, as is the case in resting cells, CaM_N_ disengages from *p*eEF-2K_TR_.

As noted earlier, the CaM_C_ metal-binding sites also appear to be occupied by Ca^2+^ in the CaM•*p*eEF-2K_TR_ complex, contrasting our previous studies on the CaM•CTM-pep^31^ and CaM•eEF-2K_TR_^36^ complexes, both carried out in the absence of Mg^2+^, using high- and low-resolution NMR approaches, respectively. Given the similar open configuration of CaM_C_ in the complexes with *p*eEF-2K_TR_ and CTM-pep, it is apparent that minor rearrangements of its Ca^2+^-binding EF-hands would suffice to accommodate Ca^2+^. Indeed, a difference in the engagement of CaM_C_ by the CTM (a shift of approximately half a helical turn) is seen upon comparing the two cases; the small accompanying changes in the corresponding EF-hands appear sufficient to accommodate Ca^2+^ (Extended Data Fig. 8).

It has been shown that the maximal activity of eEF-2K towards a peptide substrate is independent of Ca^2+^ with similar *k*_cat_ values obtained in its absence (11 sec^−1^) or presence (25 sec^− 1^), contrasting a ~900-fold increase in the apparent Ca^2+^-induced CaM-affinity (37 μM versus 42 nM)^37^. The similar *k*_cat_ values suggest that divalent cations (Mg^2+^ or Ca^2+^) within CaM minimally influence the nature of the active complex. Instead, the predominant role of these ions is to modulate the overall affinity of the CaM/eEF-2K_TR_ interactions, i.e., to alter the “concentration” of the complex through additional interactions largely independent of those that define the active state.

### Coupling of CaM_C_ to the kinase active-site

The loop (regulatory element, RE) that connects the eEF-2K_TR_ CTM and KD contains a ^<u>96</u>^PDPWA^<u>100</u>^ sequence, fully conserved in vertebrates (^<u>97</u>^DPW^<u>99</u>^ is universally conserved), that has been deemed essential for activity^38^. In the CaM•*p*eEF-2K_TR_ complex, D97, aligned through the flanking P96 and P98, hydrogen-bonds to the main-chains of W99 and A100 using its side-chain and main-chain, respectively. This conformation generates a bulge within the RE, placing W99 within a pocket created by the side-chains of P98, F102, F138, R148, V168, and Y231 (Fig. 4a). This configuration aligns a “spine” (invoking Kornev et al.^39^) comprising of CaM_C_ and highly-conserved, predominantly hydrophobic (except R148) residues of the CTM, RE, and KD_N_ to support the energetic coupling of CaM-binding to the active site through the bound ATP (Fig. 4b).

**Fig. 4.**
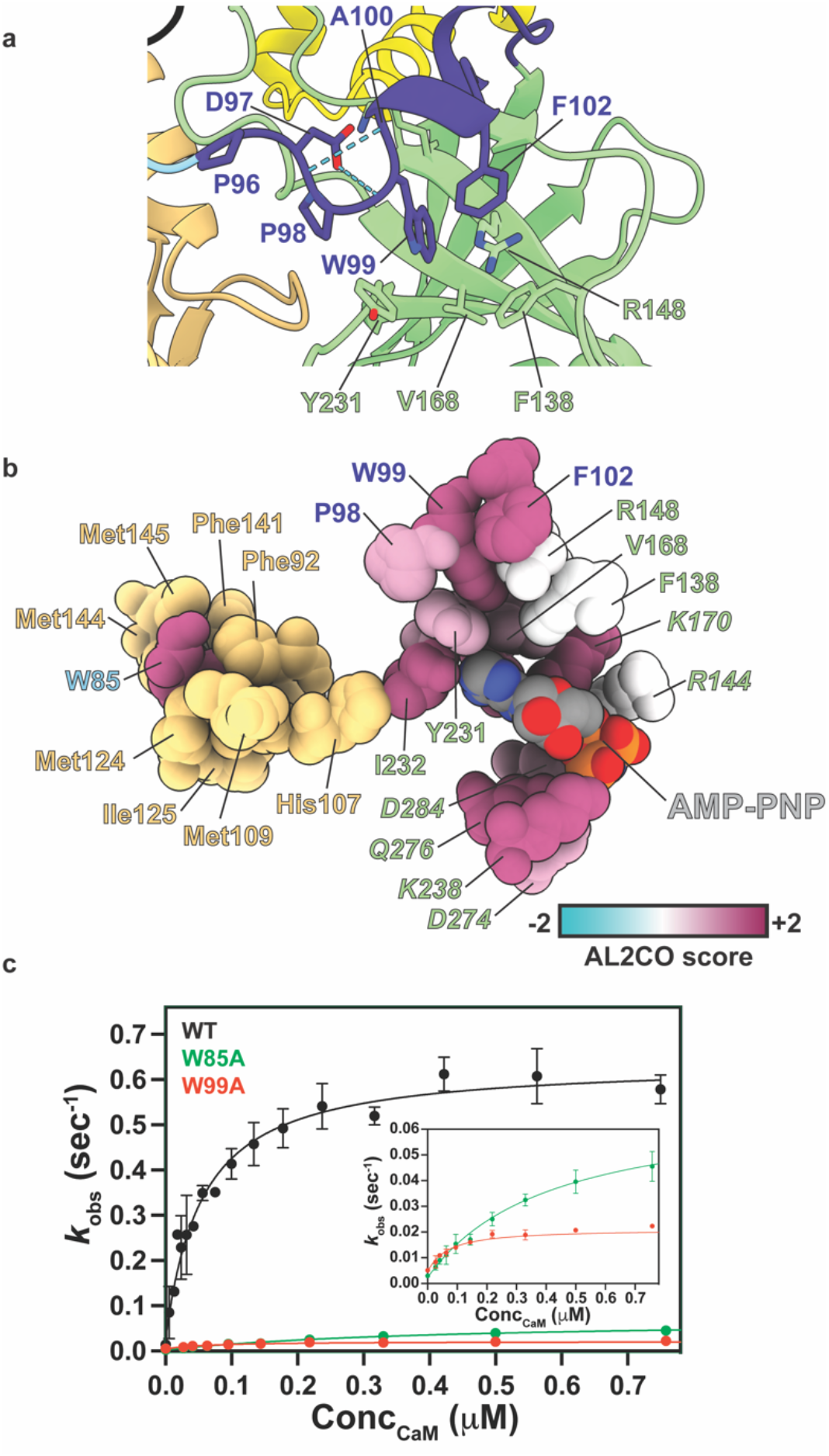
Activation of eEF-2K_TR_ by CaM. **a**, Interactions of the RE with KD_N_. **b**, Residues of the spine connecting CaM_C_ (yellow) to the eEF-2K_TR_ active site. eEF-2K_TR_ residues are colored according to conservation scores (cyan: variable, maroon: fully-conserved). AMP-PNP has been modeled in. **c**, Activity of wild-type eEF-2K (WT, black), and corresponding W85A (green) and W99A (red) mutants. The maximal catalytic rates (sec^−1^) are WT:0.61±0.02, W85A:0.07±0.002, W99A:0.02±0.001; the corresponding affinities (nM) are 52±8, 406 ± 47, 62 ± 19. The inset shows an expansion for the two mutants. Error bars indicate standard deviations over *n*=2 (WT) and *n*=3 (mutants) measurements.

Given the close-packed nature of the spine in the CaM•*p*eEF-2K_TR_ complex, one may predict that its disruption would have profound functional consequences. For example, a W99A mutation is expected to be substantially more perturbative than replacement by a similarly sized Leu, as borne out by functional studies^40^. Further, P98 hydroxylation^38^ or a His107Lys mutation^20^, both of which would significantly misalign the spine, have been shown to be deleterious to function. This effect contrasts a more modest impact of a less disruptive His107Ala mutation^20^. Additionally, sequence analyses suggest that mutations in the least, but still significantly conserved, spine residues, F138 and R148, co-vary with a nearby catalytic residue, R144. These are P139/P149/V145 in *Ursus americanus* and V137/V147/K143 in *Bos mutus*, perhaps resulting in non-functional enzymes. Thus, in our proposed model, W85 provides intrinsic binding energy^41^ for the interaction with CaM_C_31, which is utilized, in part, to stabilize the configuration of the spine and concomitantly the active state of the kinase; W99 only contributes to the latter effect. This mechanism is consistent with the observation that a W85A mutation leads to an ~9-fold reduction in the catalytic activity and an ~8-fold decrease in the CaM-affinity; a W99A mutation leads to a ~30-fold decrease in the catalytic activity without affecting CaM-affinity (Fig. 4c).

### Conclusions and outlook

Our structure of the CaM•*p*eEF-2K_TR_ complex provides a framework to interpret the large body of biochemical and functional data generated by almost four decades of work on this unique enzyme in atomic detail. This structure, combined with current and previous biochemical/biophysical measurements, suggests a central role of CaM_C_ in activating eEF-2K and defining the nature of the active state. In contrast, CaM_N_ has a more supporting role, serving primarily as a sensor of dynamic changes in Ca^2+^, e.g., during neuronal Ca^2+^ spikes/waves, in the face of a relatively constant Mg^2+^ level, to enhance the cellular concentration of active complex. This regulation ensures maintenance of basal levels of the active species and optimal elongation rates through eEF-2 phosphorylation under normal cellular conditions while allowing its rapid modulation in response to specific signals.

However, the current structure leaves several unresolved questions about eEF-2K regulation. One such puzzle concerns the mechanism of the stimulatory auto-phosphorylation on S500^42^, which is also a target of protein kinase A (PKA)^43^. S500 is buried in a shallow pocket lined by the side-chains of H260, E264, H268, and F309 on the face opposite key catalytic elements, more than 20 Å away from the base D274 (Extended Data Fig. 9a). Auto-phosphorylation *in cis* could, in principle, be achieved by a partial unfolding of α1’ with the added flexibility provided by an intact R-loop in full-length eEF-2K. Indeed, auto-phosphorylation on S500 is extremely slow and requires prior phosphorylation on T348^37^; S500 phosphorylation is not detected in *p*eEF-2K_TR_. Alternatively, auto-phosphorylation *in trans* (or phosphorylation by PKA) would require less substantial structural transitions and occur within a transiently assembled 2:2 species with an anti-parallel arrangement of its constituent CaM•*p*eEF-2K_TR_ subunits. Such an assembly would be compatible with our previous crosslinking MS (XLMS) results^22^, which are not consistent with the current heterodimeric structure (Extended Data Fig. 9b), without invoking dramatic conformational changes. Native MS studies on the CaM•eEF-2K_TR_ complex reveal charge states corresponding to a 2:2 complex, albeit at extremely low levels^22^. Similar questions regarding the conformational changes that enable (or follow) other regulatory phosphorylation events, e.g., on S366 (a target for p90^RSK44^), also remain unresolved since several of these modulatory R-loop sites are missing in eEF-2K_TR_.

## Acknowledgments

The authors thank Mr. Fatlum Hajredini and Dr. Kwangwoon Lee (Harvard Medical School) for their comments about the work. Dr. Kevin Battaile (New York Structural Biology Center) is thanked for his support during data collection and analysis.

## Author Contributions

AP prepared samples for all biophysical measurements and optimized crystallization conditions together with EAI; AP and EAI collected crystallographic data; AP calculated and refined the structural model with support from EAI and DJ; AP performed and analyzed all the solution NMR measurements; KL, AB, and EAK performed the functional studies; NW designed the eEF-2K_TR_ construct; KND and RG developed the project; AP generated the first draft of the manuscript and figures that KND and RG subsequently refined with input from all the authors.

## Data availability

Atomic coordinates for the CaM•*p*eEF-2K_TR_ complex have been deposited in the Protein Data Bank (PDB) with accession code 7SHQ.

## Methods

### Crystallization of the CaM•pTR complex

The protocols used for the expression and purification of eEF-2K_TR_ and CaM, the selective phosphorylation of the former at T348 (to generate *p*eEF-2K_TR_), and the subsequent purification of the 1:1 heterodimeric CaM•*p*eEF-2K_TR_ complex were performed as described previously^22,30^. The final samples used in the crystallization trials consisted of ~11.0 mg/mL of the CaM•*p*eEF-2K_TR_ complex in a buffer containing 20 mM Tris (pH 7.5), 100 mM NaCl, 3.0 mM CaCl_2_, 1.0 mM tris(2-carboxyethyl)phosphine (TCEP, GoldBio), 1.5 mM MgCl_2_ and 1.0 mM AMP-PNP (Millipore Sigma). Initial screens to identify crystallization conditions were performed under oil at the National Crystallization Center at the Hauptman-Woodward Medical Research Institute. Potential conditions were further optimized in-house, and optimal conditions were identified as the following: 100 mM Bis-Tris (pH 6.9), 200-300 mM MgCl_2_, and 20-26% PEG3350 combined in a 1:1 or 2:1 ratio plated on Greiner 72-well micro-batch plates (Hampton Research) under paraffin oil (EMD Chemicals) at ambient temperature. Multiple crystal clusters emerged in less than 12 hours, reaching their maximum size within a few days. Most of these, however, were extremely fragile and diffracted poorly. A single condition, 1:1 ratio, 24% PEG3350, and 300 mM MgCl_2_, yielded crystals that produced diffraction data of good quality.

### Structure determination

Data were collected at the NSLS-II light source at Brookhaven National Labs utilizing the 19-ID (NYX) beamline. Two datasets collected on distinct regions of the same crystal were processed and combined using the autoPROC toolbox^45^ resulting in a dataset with a resolution of 2.3 Å. The crystal (P3_1_21) with an elongated unit cell (a = b = 58.49 Å, c = 365.78 Å) contained a single CaM•*p*eEF-2K_TR_ heterodimer in the asymmetric unit. Initial phases were obtained using phemix.mr_rosetta^46^ and an alignment file obtained from the HHpred server^47^ containing 53 entries, including sequences corresponding to the crystal structures of the α-kinase domains of MHCK-II^25^, ChaK^23^, together with those of several SEL-1 and TPR repeat-containing proteins. The initial model was built using multiple cycles consisting of Phaser^48^, Coot^49^, and phenix.refine^50^. At the end of the initial build, further optimization was carried out using the PDB_REDO server^51^, yielding a model that contained 469 eEF-2K_TR_ residues (88%) and the C-lobe of CaM. Most of the CaM N-lobe (except the loop between helices α_C_ and α_D_) was built through iterative manual fitting with Coot followed by refinement using Phenix. A comparison of the CaM N-lobe fold with existing CaM structures using the DALI server^52^ showed its close resemblance to a Mg^2+^-bound closed form. Indeed, molecular replacement using Phaser and PDB:3UCW^33^ together with the rest of the CaM•eEF-2K_TR_ complex structure yielded a nearly complete model. The remaining missing segments of the structure of the complex were then progressively added and improved using Coot, phenix.refine, and ISoLDE^53^. Cations were identified using the CheckMyBlob server^54^ and inserted with solvent molecules using Coot and refined using Phenix. The final structural model consisted of 493 and 145 residues of eEF-2K_TR_ (92.8%) and CaM (97.9%), respectively, together with 1 Zn^2+^, 2 Ca^2+^, 5 Mg^2+^ ions and 145 water molecules. No discernible density for AMPPNP was observed. Details about data collection and refinement are shown in Extended Data Table 1.

The sequence conservation in eEF-2K_TR_ was obtained using the ConSurf server^55^ with default parameters that generated a comparison with 150 non-redundant sequences (95% identity cut-off). The generated alignment file was read into UCSF ChimeraX^56^ and represented on the eEF-2K_TR_ structure with the corresponding residues colored by their AL2CO scores^57^.

### Solution NMR spectroscopy

^13^C,^1^H-Ile-δ1, Met-*ε*, ^2^H,^15^N-labeled CaM (IM-labeled CaM)^36^ and unlabeled *p*eEF-2K_TR_30 were prepared as previously described, and the corresponding 1:1 heterodimeric complex was purified as described above. All NMR samples (unless specifically noted) were prepared in a buffer (NMR buffer) containing 20 mM HEPES (pH 7.5), 100 mM NaCl, 1.0 mM TCEP, and 5% ^2^H_2_O. The following samples were prepared: (1) IM-labeled CaM alone (~100 μM) in NMR buffer containing 3.0 mM CaCl_2_ or (2) as part of the corresponding CaM•*p*eEF-2K_TR_ complex (~147μM); (3) IM-labeled CaM (~100 μM) in NMR buffer containing 5.0 mM EGTA and 310 mM MgCl_2_; (4) IM-labeled CaM alone (~100 μM) in NMR buffer containing 3.0 mM CaCl_2_ and 300 mM MgCl_2_ or (5) as part of the corresponding CaM•*p*eEF-2K_TR_ complex (~147 μM); (6) ~50 μM samples of IM-labeled CaM in NMR buffer containing ~150 μM DSS and 0, 0.2, 0.4, 5.0, 20, 103 or 310 mM MgCl_2_ (all samples contained EGTA in a ~1/60 ratio with respect to Mg^2+^); (7) samples containing ~18 μM of the CaM•*p*eEF-2K_TR_ complex in buffer comprising 20 mM HEPES (pH 7.5), 10 mM NaCl, 0.5 mM TCEP, ~150 μM DSS, and 0.3 mM CaCl_2_ with 0, 1.0, 3.0, 6.0, 12.0, 36, 68, 115, 240 and 450 mM MgCl_2_. ^1^H,^13^C SOFAST-HMQC^58^ experiments were carried out on all samples using sweep-widths of 13.95 ppm (512 complex points) and 12.0 ppm (128 complex points) in the direct and indirect dimensions, respectively. All experiments were carried out at 25 °C on an 800 MHz Bruker Avance-III HD spectrometer equipped with a triple-resonance cryogenic probe capable of applying pulsed-field gradients along the z-axis. Data were processed using nmrPipe^59^ and analyzed using nmrViewJ^60^. Chemical shift perturbations (Δδ in ppm) were calculated using^36^

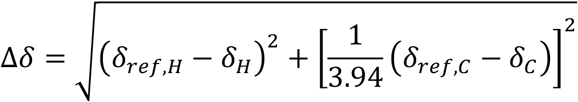

Where δ_*ref,H*_ and δ_*ref,C*_ are the reference ^1^H and ^13^C methyl chemical shifts, respectively, and δH and δC are the corresponding shifts in the presence of relevant additives.

### Small-angle X-ray scattering

Small-angle X-ray scattering (SAXS) data were acquired on two samples containing 1.9 mg/mL (lc) and 6.4 mg/mL (hc) of the CaM•eEF-2K_TR_ complex (unphosphorylated eEF-2K_TR_), as previously described^22^. These data were reanalyzed using ATSAS 3.0.1 software^61^, and identical results as those shown in Table 3 of Will et al.^22^, were obtained. However, a slightly different approach was utilized to obtain *ab initio* 3-dimensional models from the SAXS data. 25 structures were calculated for each of the hc and lc datasets using DAMMIF^62^ using default parameters in the slow mode and clustered using DAMCLUST^63^. For the hc dataset, a total of 6 clusters were obtained in which 3 (hc_1, hc_2, hc_4) contained a single member, 1 each contained 2 (hc_5) or 3 (hc_3) members, and the largest cluster (hc_6; NSD to all other clusters = 1.47±0.31) comprised of 17 members. The averaged envelope from 6_hc was refined against the experimental hc dataset (χ^2^=0.9766) using DAMMIN^64^ (run in the slow mode using default parameters) by restricting the search volume using DAMSTART. For the lc dataset, 2 clusters contained a single member (lc_1, lc_2), 1 each contained 3 (lc_5), 5 (lc_6), 7 (lc_3) or 8 members (lc_4). Thus, the averaged structures from the two largest clusters (lc_3 and lc_4, NSD=1.16) were separately refined against the experimental lc dataset (χ^2^=0.7761 and χ^2^=0.7767 for lc_3 and lc_4, respectively).

The SAXS profile was calculated from the structure of the CaM•*p*eEF-2K_TR_ complex using CRYSOL3^65^ and fitted separately to the lc and hc datasets yielding χ^2^ values of 1.504 and 3.217, respectively, suggesting that the former provides a somewhat better agreement with that expected from the structure. The molecular envelopes calculated from each dataset (lc_3, lc_4, hc_6) were superimposed on the crystal structure using SUPCOMB^66^. The structure of the CaM•*p*eEF-2K_TR_ complex fitted into the refined molecular envelope of the lc_3 cluster and a comparison of the theoretical and calculated data for the lc dataset are shown in Extended Data Fig. 2.

### Measurement of eEF-2K_TR_ activity in the presence of Mg^2+^

Auto-phosphorylation of eEF-2K_TR_ (500 nM) was carried out in buffer comprising 25 mM HEPES (pH 7.0), 2 mM DTT, 40 μg/mL BSA, 50 mM KCl, and 5 μM CaM. The reaction was conducted under two distinct conditions – one with 15 mM MgCl_2_ and 0.15 mM CaCl_2_ and a second with 150 mM MgCl_2_ and 1.5 mM CaCl_2_. The reaction mixture was incubated at 30 °C for 10 min, and the reaction was initiated by the addition of 500 μM [γ-^32^P]ATP (100−1000 c.p.m./pmol) in a final volume of 250 μL. Aliquots (10 pmol/20 μL) of eEF-2K_TR_ were removed every 0, 10, 23, 32, 40, 60, 120 and 300 s over a 5 min time period and directly added to hot SDS−PAGE sample loading buffer containing 125 mM Tris-HCl (pH 6.75), 20% glycerol (v/v), 10% 2-mercaptoethanol (v/v), 4% SDS, and 0.02% bromophenol blue) to quench the reaction. The mixture was further heated for 5 min at 95 °C to ensure complete denaturation. The samples were resolved by SDS−PAGE and stained with Coomassie Brilliant Blue. Gels were exposed for 8 h in a phosphorimager cassette, scanned in a Typhoon 9500 imager, and analyzed using ImageJ. Autophosphorylation was quantified by drying the gels and excising the eEF-2K_TR_-containing segments. The radioactivity was measured with a Packard 1500 liquid scintillation analyzer.

### Measurement of the activity of full-length eEF-2K and mutants

Assays were performed in buffer containing 25 mM HEPES, 50 mM KCl, 10 mM MgCl_2_, 150 μM CaCl_2_, 20 μg/mL BSA, 100 μM EGTA, 2 mM DTT, 0-4 μM CaM, and 10 μM sox-peptide^67^. The assays used 5 nM, 10.5 nM, or 8.6 nM of unphosphorylated wild-type eEF-2K, the W85A mutant, or the W99A mutant, respectively. The reaction was initiated with the addition of 1 mM ATP. Product turnover was monitored by fluorescence (excitation at 360 nm, emission at 482 nm) using a Synergy H4 plate reader (BioTek). The data were analyzed using a quadratic binding isotherm to obtain *k*_obs,max_, and *K*_CaM_ values. Experiments were performed in duplicate (for wild-type) or in triplicate (for the W85A and W99A mutants).

## Extended Data and Tables

**Extended Data Fig. 1.**
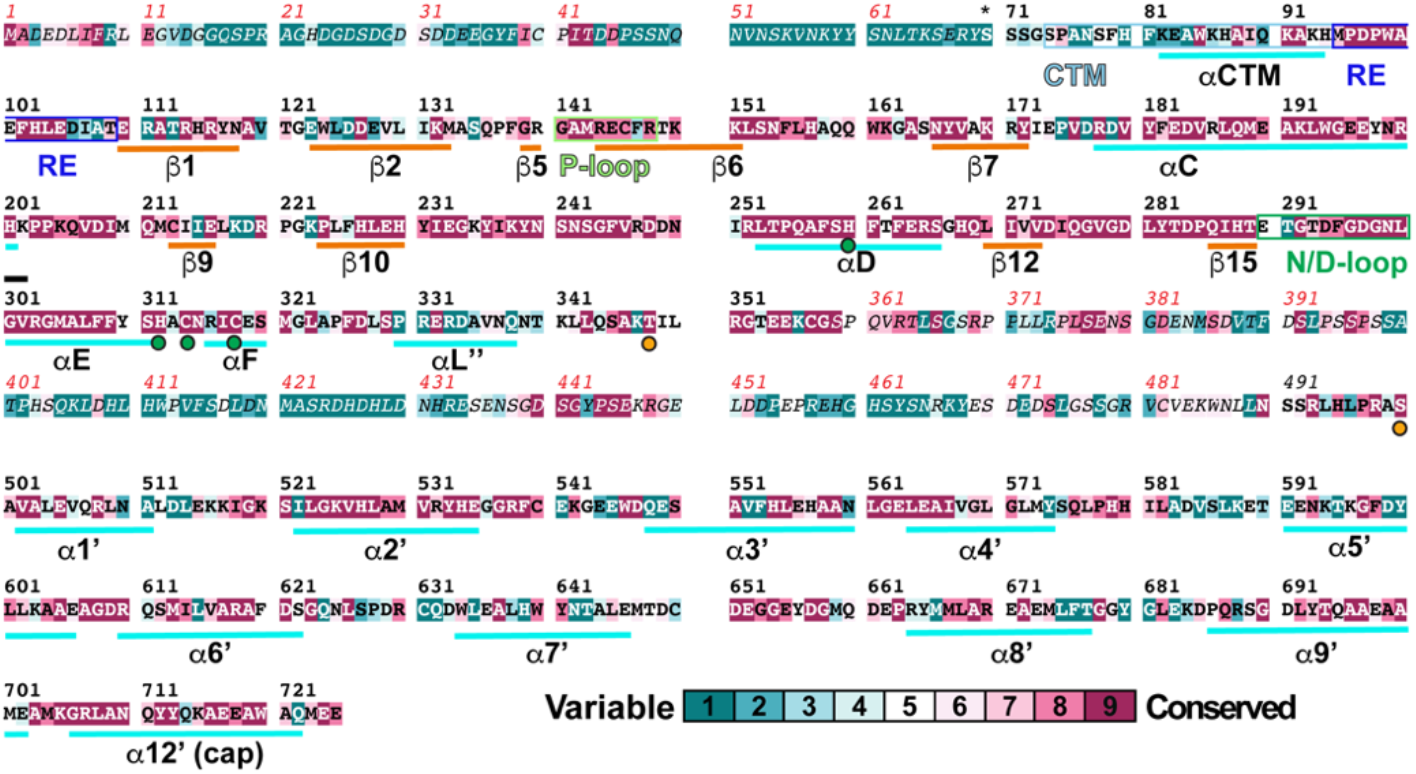
Sequence conservation in eEF-2K. Sequence conservation in eEF-2K was obtained using the ConSurf server with standard settings. Specific regions of secondary structure as determined by UCSF ChimeraX are indicated below the sequence; the location of the CaM-targeting motif (CTM, 74-95; previously known as the CaM-binding domain, CBD), regulatory element (RE, 96-109), P-loop (141-148) and N/D-loop (290-300, previously termed the G-loop), are also indicated. Labeling of the secondary structural elements for the KD (110-321) is based on that previously assigned to myosin heavy chain kinase A (MHCK-A). Green circles indicate residues H260, H312, C314, and C318 that coordinate the structural Zn^2+^ found in all α-kinases. The orange circles indicate the primary (T348) and secondary (S500) auto-phosphorylation sites. The secondary structural elements for the C-terminal domain (CTD, 502-725) are primed. The 359-489 segment of the R-loop (321-501) has been replaced by a 6-glycine linker in eEF-2K_TR_ (previously called TR) that is also missing the first 69 residues of full-length eEF-2K; S70 (indicated by a ‘*’) that is native in eEF-2K re-appears as a remnant of cloning. The residues present or absent in the eEF-2K_TR_ construct compared to full-length eEF-2K are shown in bold or italicized fonts, respectively, with the corresponding sequencing numbering indicated in black or red font.

**Extended Data Fig. 2.**
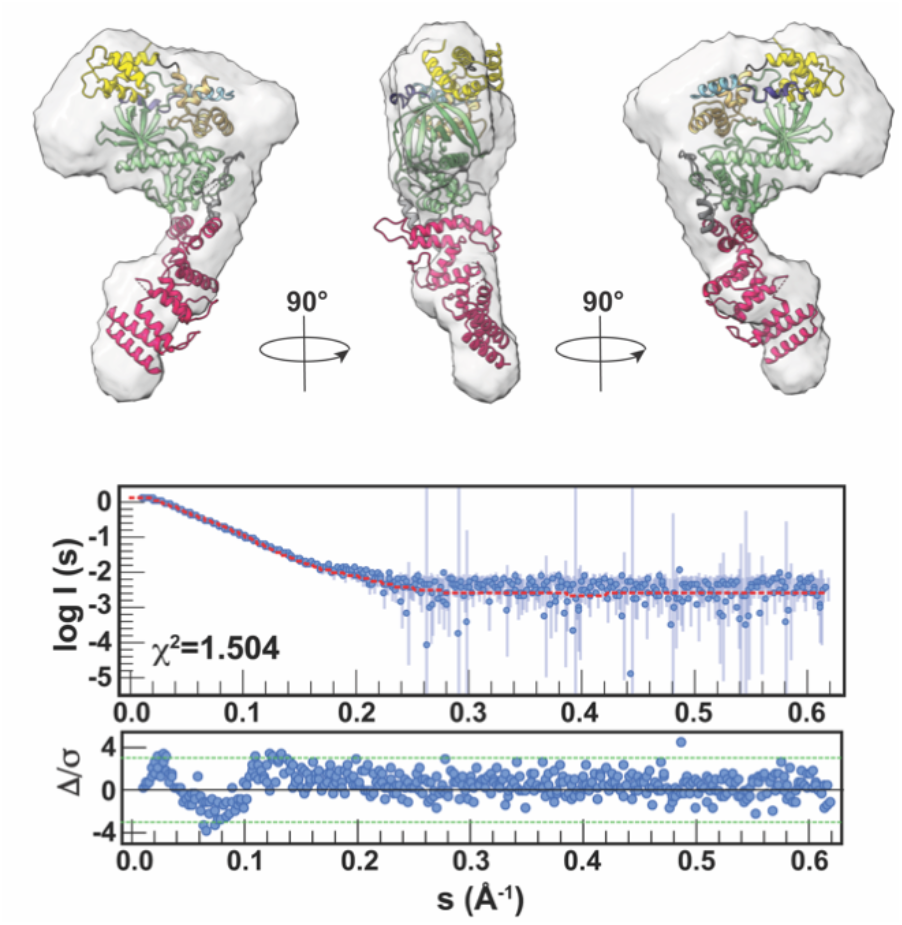
The structure of the CaM•*p*eEF-2K_TR_ complex is consistent with solution scattering data. The structure of the CaM•*p*eEF-2K_TR_ complex fitted into the molecular envelope of one of two dominant clusters calculated from small-angle X-ray scattering (SAXS) data acquired on a low-concentration sample of the CaM•eEF-2K_TR_ complex (unphosphorylated T348). The corresponding experimental data (blue circles), the theoretical fit (red curve), and the reduced residuals (Δ/σ) are shown on the lower panel. While the SAXS-generated envelope reproduces the overall structural features in solution, the reduced residuals at low *s* values, and the relatively large χ^2^ value, suggest differences between the molecular species. Some of these differences can be attributed to the fact that the SAXS data were acquired on the unphosphorylated complex in the presence of Ca^2+^ but in the absence of Mg^2+^.

**Extended Data Fig. 3.**
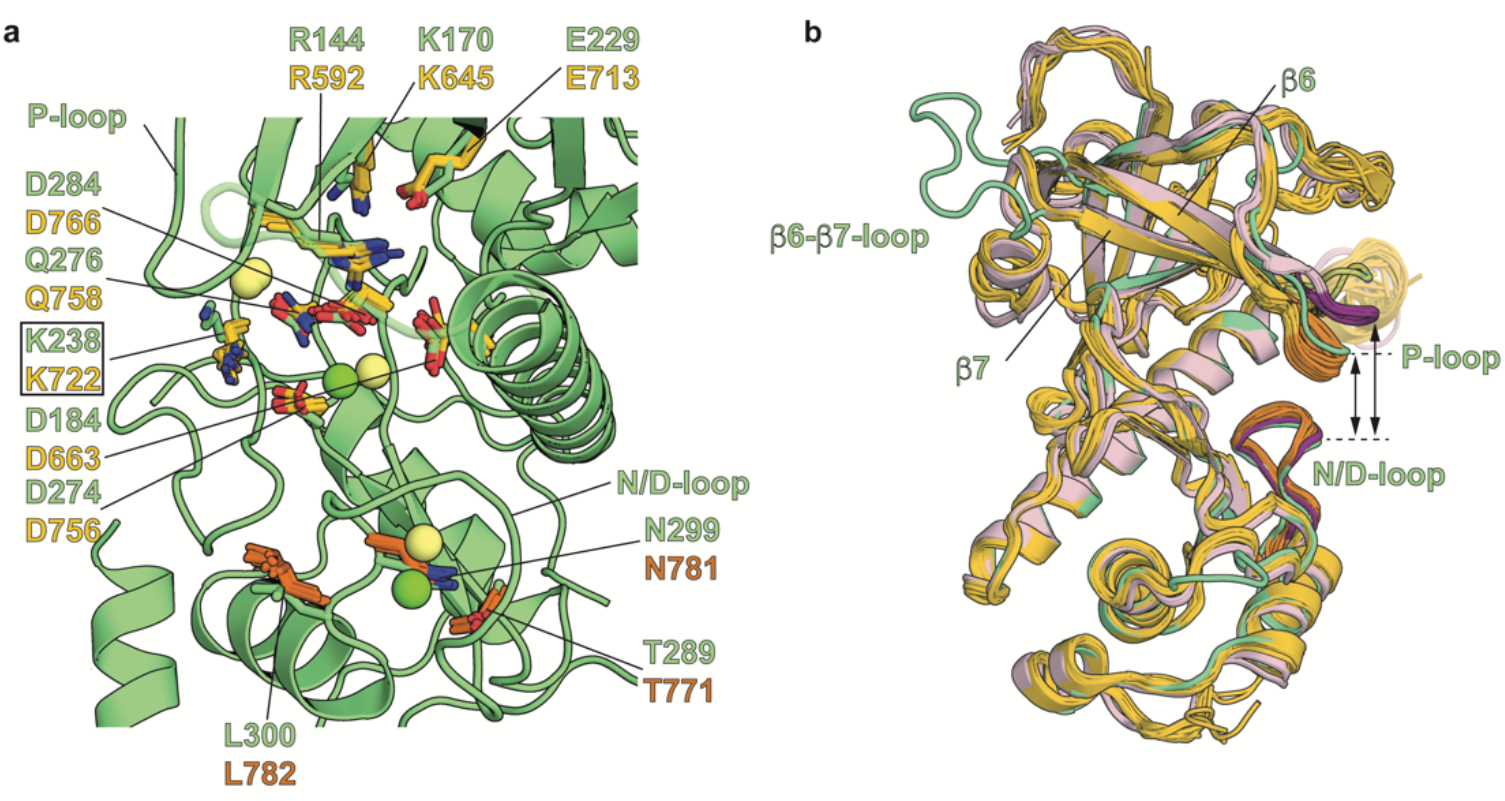
*p*eEF-2K_TR_ is in an active state in the CaM•*p*eEF-2K_TR_ complex. **a**, Comparison of the orientations of key “catalytic” residues (in green) of the eEF-2K_TR_ KD in the CaM•*p*eEF-2K_TR_ complex with corresponding residues (light orange) in several structures of the MHCK-A KD. R144 of eEF-2K_TR_ (R592 in MHCK-A) coordinates the α- and β-phosphates of ATP; K170 (K645) engages the α-phosphate and the adenine ring; E229 (E713) engages the adenine amino group; K238 (K722) engages the γ-phosphate of ATP; D274 (D756) is the putative catalytic base, Q758 (Q276) and D284 (D766) coordinate the catalytic Mg^2+^ ions and the γ- and α-phosphates of ATP, respectively. While no density corresponding to a nucleotide was seen in the CaM•*p*eEF-2K_TR_ complex (crystallized in the presence of the slowly hydrolyzable ATP analog, AMP-PNP), all side-chains (except K238, indicated by the rectangle) are in similar orientations as their nucleotide-bound MHCK-A counterparts. The Mg^2+^ ions seen in the structure of the CaM•*p*eEF-2K_TR_ complex are shown green, and those seen in the MHCK-A structures are shown in lime-green. Also shown (MHCK-A residues in orange) are the N/D-loop residues, T289 (T771), N299 (N781), and L300 (L782). N299 is stabilized through hydrogen bonds with the T289 side-chain and the D294 main-chain. This “asparagine-in” conformation has been suggested to be a characteristic of the active state of MHCK-A. In MHCK-A, N781 and L782 have been suggested to determine the accessibility of the active site. Given their similar spatial location and orientations, a similar role for the analogous N299 and L300 in eEF-2K_TR_ can be envisaged. The conserved D766 in MHCK-A was phosphorylated in some of the structures. This phospho-aspartyl species has been proposed to represent an intermediate on the pathway to substrate phosphorylation. We do not find evidence of phosphorylation on the equivalent residue (D284) in eEF-2K_TR_ in the CaM•*p*eEF-2K_TR_ complex. None of the MHCK-A structures used in the overlay contain *p*D766. **b**, Overlay of the KD of *p*eEF-2K_TR_ in the CaM• *p*eEF-2K_TR_ complex (green) and MHCK-A KDs in their nucleotide-bound (yellow) or nucleotide-free structures (pink) with the corresponding P- and the N/D-loops colored orange and magenta, respectively. A closed conformation of the P-loop relative to the N/D-loop in the presence of nucleotide is evident. This closed configuration is also seen for the eEF-2K_TR_ KD in the CaM• *p*eEF-2K_TR_ complex. Also indicated for reference is the longer β6-β7-loop of eEF-2K_TR_ that contacts CaM_N_ in the CaM•*p*eEF-2K_TR_ complex. The following MHCK-A structures were used in the analysis: for **a**, 5E9E (bound to AMP-PNP), 3LKM (bound to AMP), 3LMI (D766A mutant bound to ATP), 3LLA (bound to AMPPCP), and 4ZS4 (D756A mutant bound to ATP); for **b**, 3LLA, 3LMH (*p*D766, bound to ADP), 4ZS4, 3PDT (C-terminally truncated bound to ADP), 3LKM, 5E9E, 5DYJ (D663A mutant bound to AMP), 4ZME (*p*D766, bound to adenosine), 4ZMF (*p*D766, bound to AMP), and 5E4H (apo) in **b**. Note that the only open conformation of the P-loop corresponds to the apo-enzyme (5E4H). In these structures, D663 was found to be phosphorylated. We do not find evidence of phosphorylation on the equivalent D184 in the structure of the CaM•*p*eEF-2K_TR_ complex. Structural alignments were obtained using the DALI server.

**Extended Data Fig. 4.**
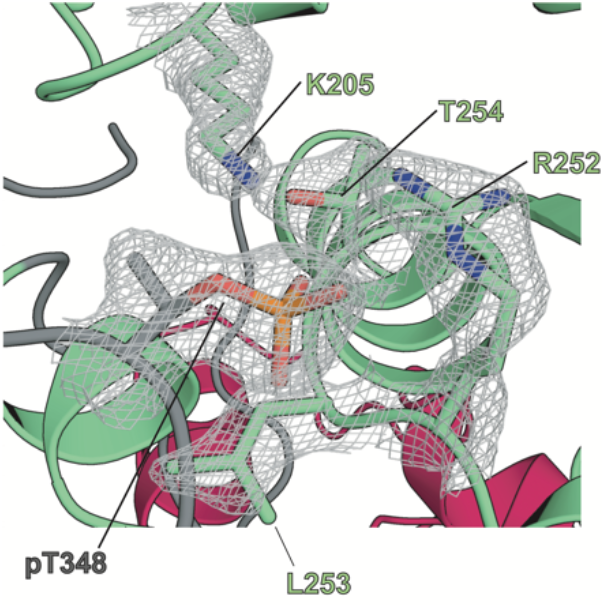
T348 is phosphorylated in the CaM•*p*eEF-2K_TR_ complex. 2Fo-Fc map contoured at 1.5σ, illustrating *p*T348 engaged at the PBP. The side-chains of key residues K205, R252, L253, and T254 are shown.

**Extended Data Fig. 5.**
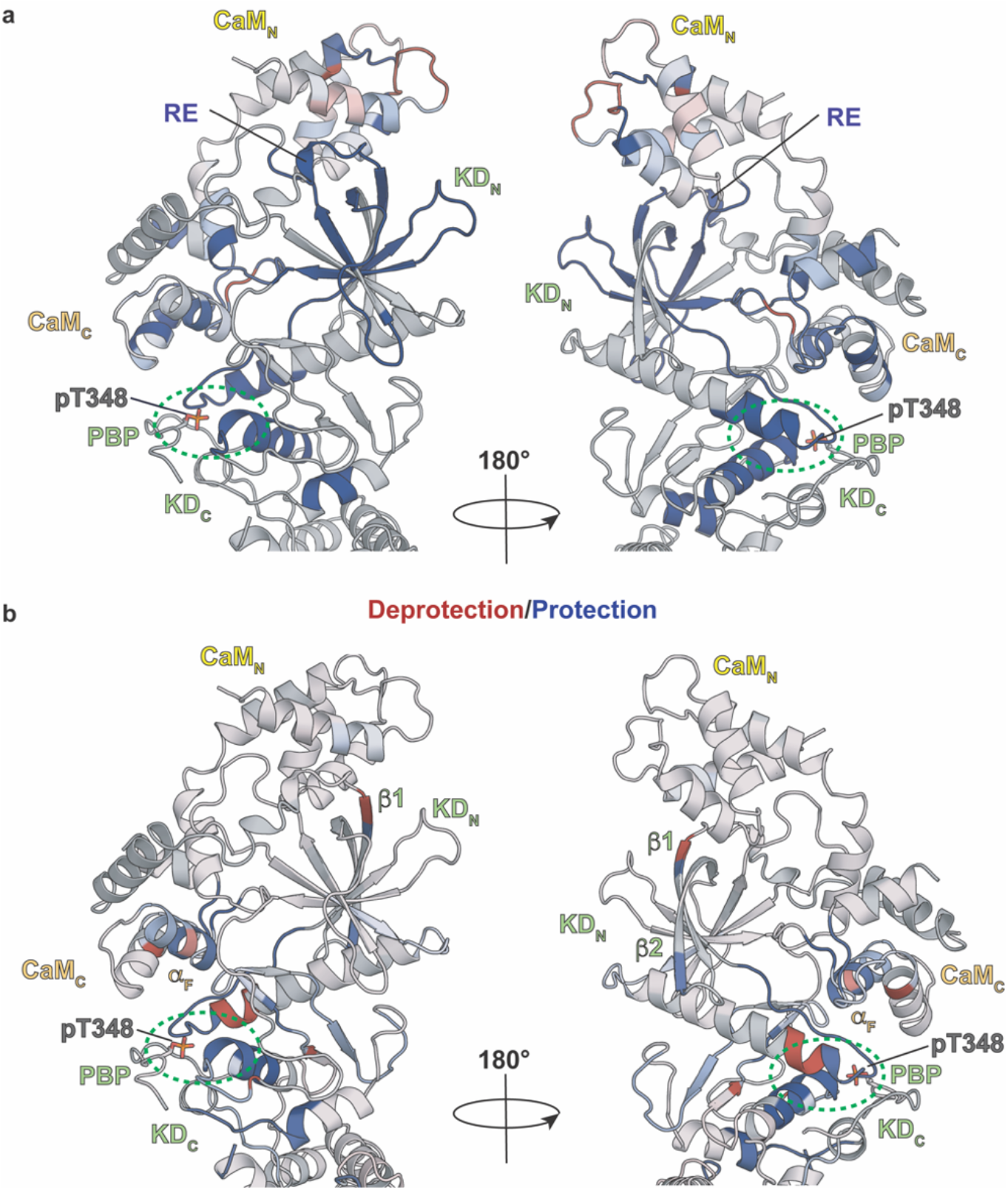
HXMS measurements reveal changes in solvent protection upon complex formation and T348 phosphorylation. **a**, Changes in solvent protection for the eEF-2K_TR_ KD and CaM upon formation of the CaM•eEF-2K_TR_ complex (unphosphorylated T348) from HXMS measurements, described previously, are plotted on the structure of the CaM•*p*eEF-2K_TR_ complex. Shades of blue or red indicate statistically significant increases or decreases in protection, respectively, upon complex formation. Increased protection is seen throughout CaM_C_, on KD_N_, including on the RE, and on KD_C_ near the PBP (indicated by the green dashed oval). This suggests the intimate coupling of the structural dynamics of CaM_C_ to key regulatory elements within the KD. Changes for CaM_N_ are heterogeneous and more modest. **b**, Changes in solvent protection upon phosphorylation on T348 comparing the CaM•eEF-2K_TR_ and CaM•*p*eEF-2K_TR_ complexes (shades of blue indicate increased protection upon phosphorylation on T348). Changes, in this case, are more local and seen at the PBP and on the proximal CaM_C_, especially on αF. Some isolated changes are also seen on KD_N_ (β1 and β2).

**Extended Data Fig. 6.**
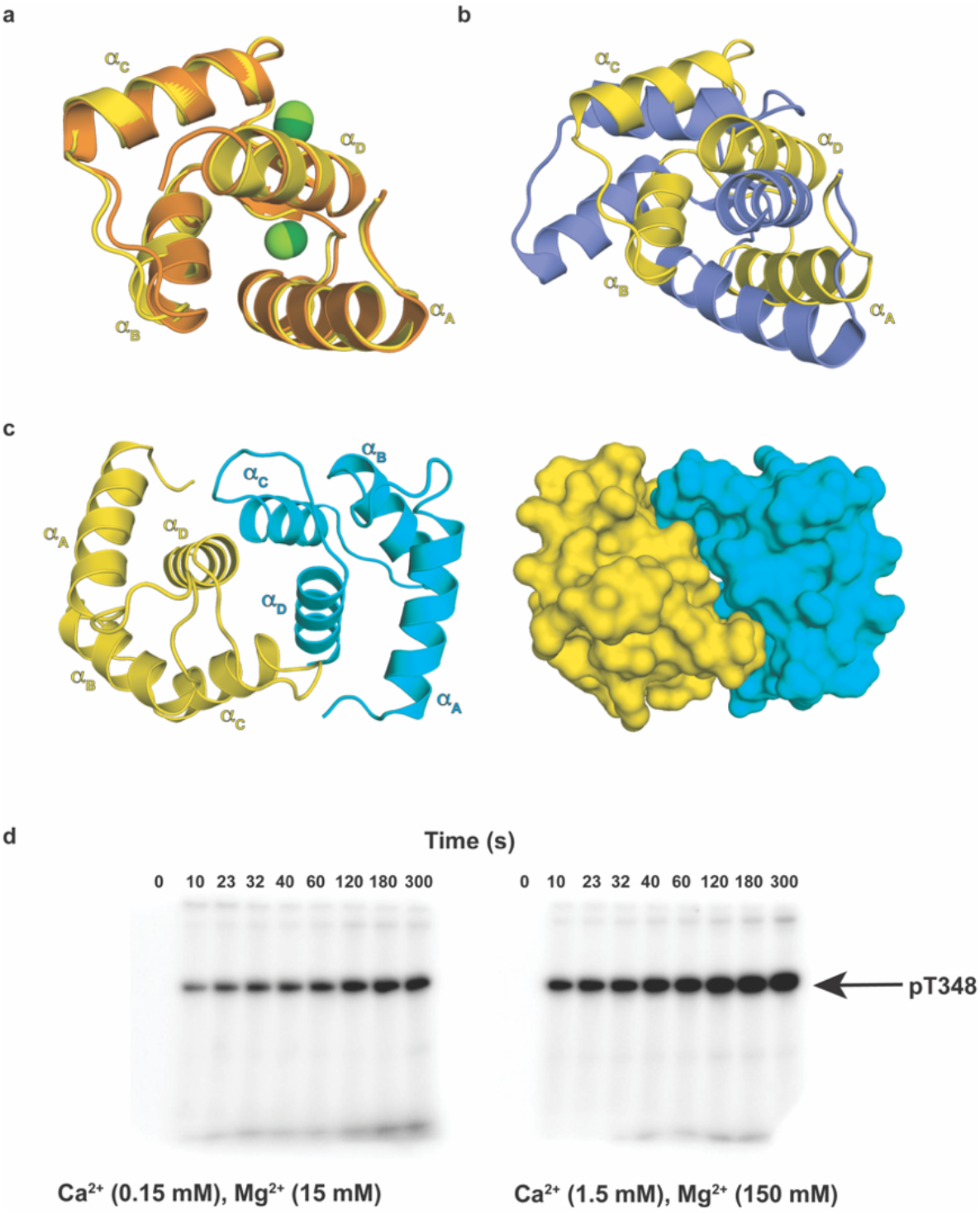
CaM_N_ adopts a closed conformation in the CaM•*p*eEF-2K_TR_ complex. **a**, Conformation of CaM_N_ in the CaM•*p*eEF-2K_TR_ complex (yellow) or that bound to Mg^2+^ (orange; PDB: 3UCW); the corresponding bound divalent cations are shown as light- or dark-green spheres. **b**, A comparison of the conformation of CaM_N_ in the CaM•*p*eEF-2K_TR_ complex (yellow) with that in the NMR structure of the complex with CTM-pep in the presence of Ca^2+^. CaM_N_ engages *p*eEF-2K_TR_ in an Mg^2+^-bound “closed” conformation rather than a Ca^2+^-bound “open” conformation as in the CTM-pep complex. The metal ions have been omitted for clarity. **c**, Interactions between two neighboring CaM_N_ units related by symmetry within the crystal are shown as ribbons (left) or surfaces (right). The extensive crystal contacts involving α_C_ and α_D_ (the interactions with *p*eEF-2K_TR_ largely involve hydrophobic residues on α_A_ and α_B_) suggest that this arrangement stabilizes the complex within the lattice. **d**, eEF-2K_TR_ retains the ability to efficiently auto-phosphorylate on T348 under conditions where the Ca^2+^ to Mg^2+^ ratio (1:100) is similar to that used for crystallization. The time-courses for T348 auto-phosphorylation measured using two different sets of concentrations for Ca^2+^ and Mg^2+^ while holding their ratio constant are shown.

**Extended Data Fig. 7.**
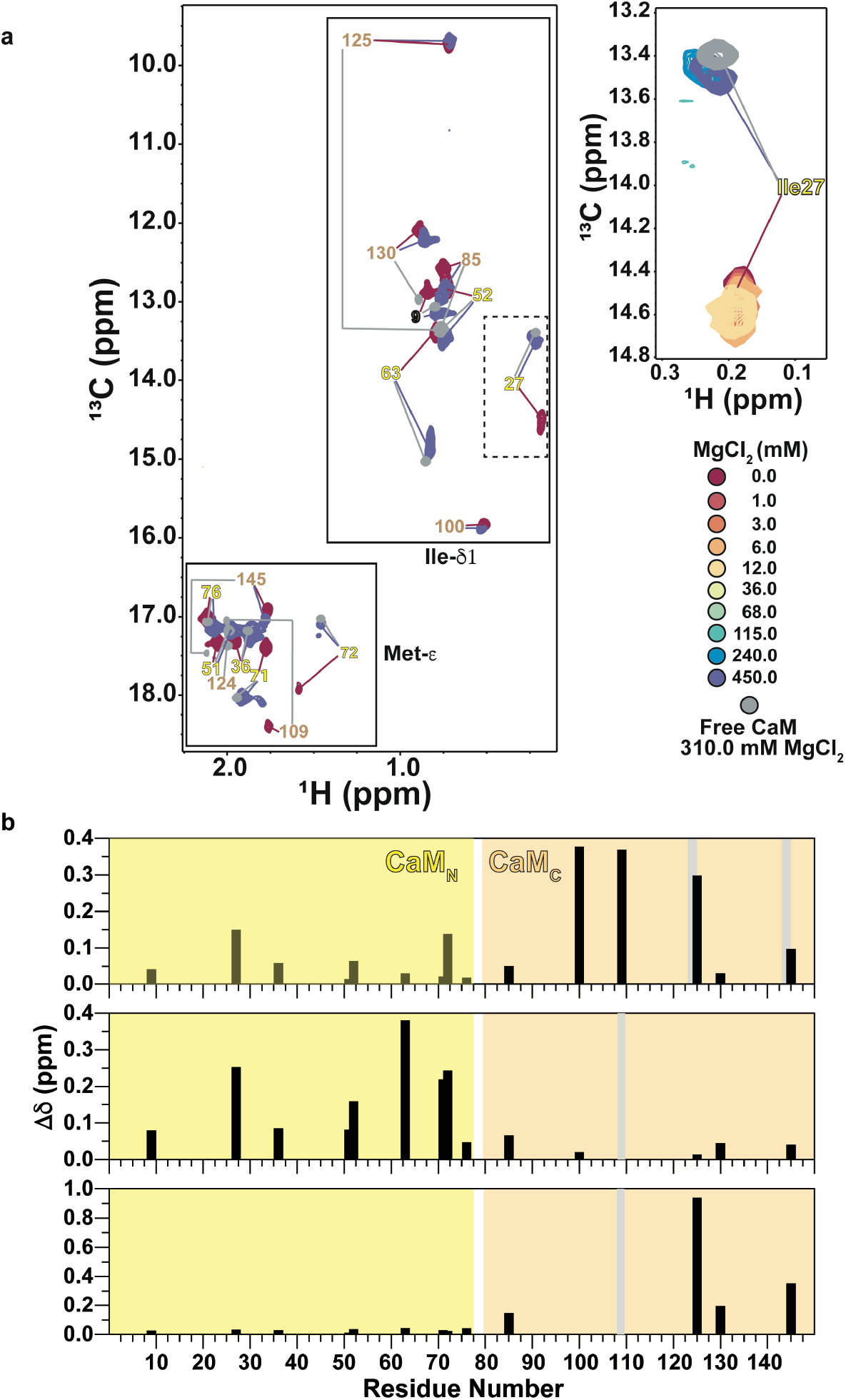
Mg^2+^ partitions exclusively to CaM_N_ in the CaM•*p*eEF-2K_TR_ complex. **a**, Overlays of ^13^C, ^1^H HMQC spectra (800 MHz) of IM-labeled CaM in the CaM•*p*eEF-2K_TR_ complex in a buffer containing CaCl_2_ (300 μM) with varying concentrations of MgCl_2_. The resonances of CaM alone in the presence of a saturating amount of MgCl_2_ are shown in grey for reference. The residues belonging to CaM_N_ or CaM_C_ are labeled in yellow and brown font, respectively. Only the initial (red, 0 mM MgCl_2_) and final (navy-blue, 450 mM MgCl_2_) points of the Mg^2+^ titration are shown on the left panel. The entire titration course for Ile27, a representative CaM_N_ resonance, is shown on the right panel. It is evident from the spectra that the CaM_N_ resonances converge to those of the Mg^2+^-bound, *p*eEF-2K_TR_-free state at very high concentrations of Mg^2+^. The perturbations experienced by the CaM_C_ resonances are significantly smaller, and they remain very close to their original positions even at 450 mM MgCl_2_. For CaM_C_, the only apparent effect is seen for Met109 (at the CTM-binding pocket), which is progressively broadened and fully disappears at the highest MgCl_2_ concentration. Multiple exchange regimes are seen for the CaM_N_ resonances. Resonances whose initial and final positions in the Mg^2+^-titration are separated by more than ~100 Hz (e.g., Ile27, Met72) are in a slow exchange regime, suggesting that Ca^2+^/Mg^2+^ exchange occurs on the ms timescale. **b**, Chemical shift perturbations induced by *p*eEF-2K_TR_ on IM-labeled CaM in the presence of Ca^2+^ (and the absence of Mg^2+^) are shown on the top panel. Fully broadened resonances are indicated by the grey bars. The largest perturbations are observed for CaM_C_. The patterns of perturbations are similar to that reported by the presence of eEF-2K_TR_ (unphosphorylated T348), suggesting that the CaM-recognition modes of eEF-2KTR and *p*eEF-2KTR are similar in the two cases. Perturbations induced by Mg^2+^ (450 mM MgCl_2_) on IM-labeled CaM in the CaM•*p*eEF-2K_TR_ complex (containing 300 μM CaCl_2_) are shown on the middle panel. Significant perturbations are seen on CaM_N_; minimal perturbations are seen for CaM_C_ (except for the Met109 resonance that is broadened to below the noise). Perturbations in IM-labeled CaM comparing the CaM•*p*eEF-2K_TR_ complex in buffer containing 300 μM CaCl_2_ and 450 mM MgCl_2_, with CaM alone in the presence of 310 mM MgCl_2_, are shown on the bottom panel. The data suggest that, at high concentrations, Mg^2+^ replaces Ca^2+^ on CaM_N_, leading to its disengagement from *p*eEF-2K_TR_.

**Extended Data Fig. 8.**
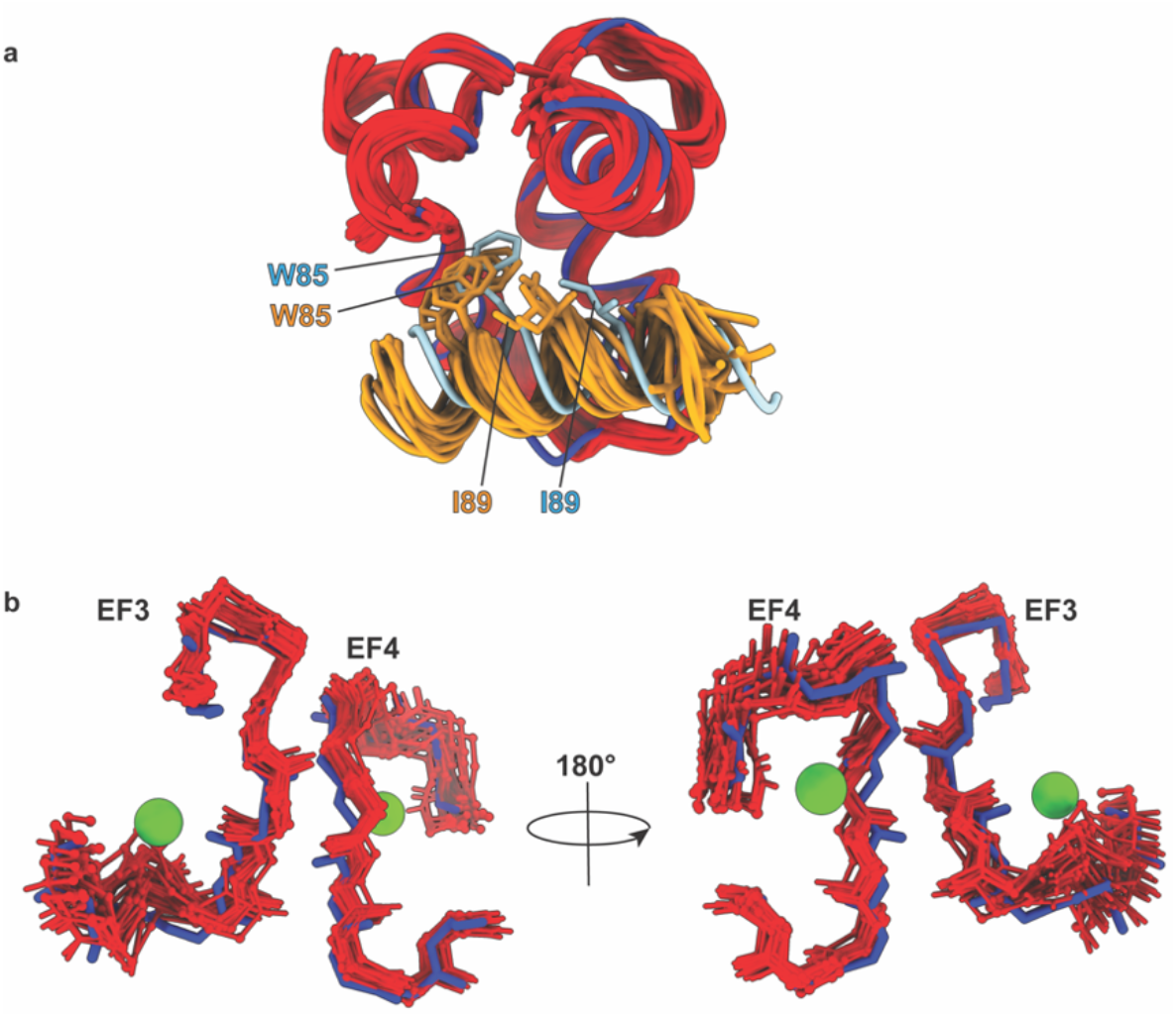
The CaM_C_/CTM module adopts slightly different conformations in the *p*eEF-2K_TR_ and CTM-pep complexes. **a**, Comparison of the CaM_C_/CTM modules in the NMR ensemble of the complex of CaM with CTM-pep (PDB: 5J8H) or in the CaM•*p*eEF-2K_TR_ complex. CaM_C_ (CTM) is shown in red (orange) and dark-blue (light-blue) in the NMR and crystal structures, respectively. The CTM is displaced by approximately half a helical turn within CaM_C_ in the crystal structure compared to the NMR ensemble. The W85 and I89 side-chains are shown for only 5 representative structures of the NMR ensemble for ease in visualization. The W85 side-chain occupies a similar spatial position, on average, in the crystal structure and the NMR ensemble. The orientation of the side-chains of I89 and A92 (hidden) are significantly different in the two cases. **b**, Backbone traces of the Ca^2+^ binding EF-hands (EF3, EF4) of CaM_C_ in the NMR ensemble (red) of the CaM•CTM-pep complex illustrate their displacement from the spatial position seen in the crystal structure of the CaM•*p*eEF-2K_TR_ complex (blue). These displacements make the CaM_C_ EF-hands unsuitable for coordinating Ca^2+^ in the CaM•CTM-pep complex.

**Extended Data Fig. 9.**
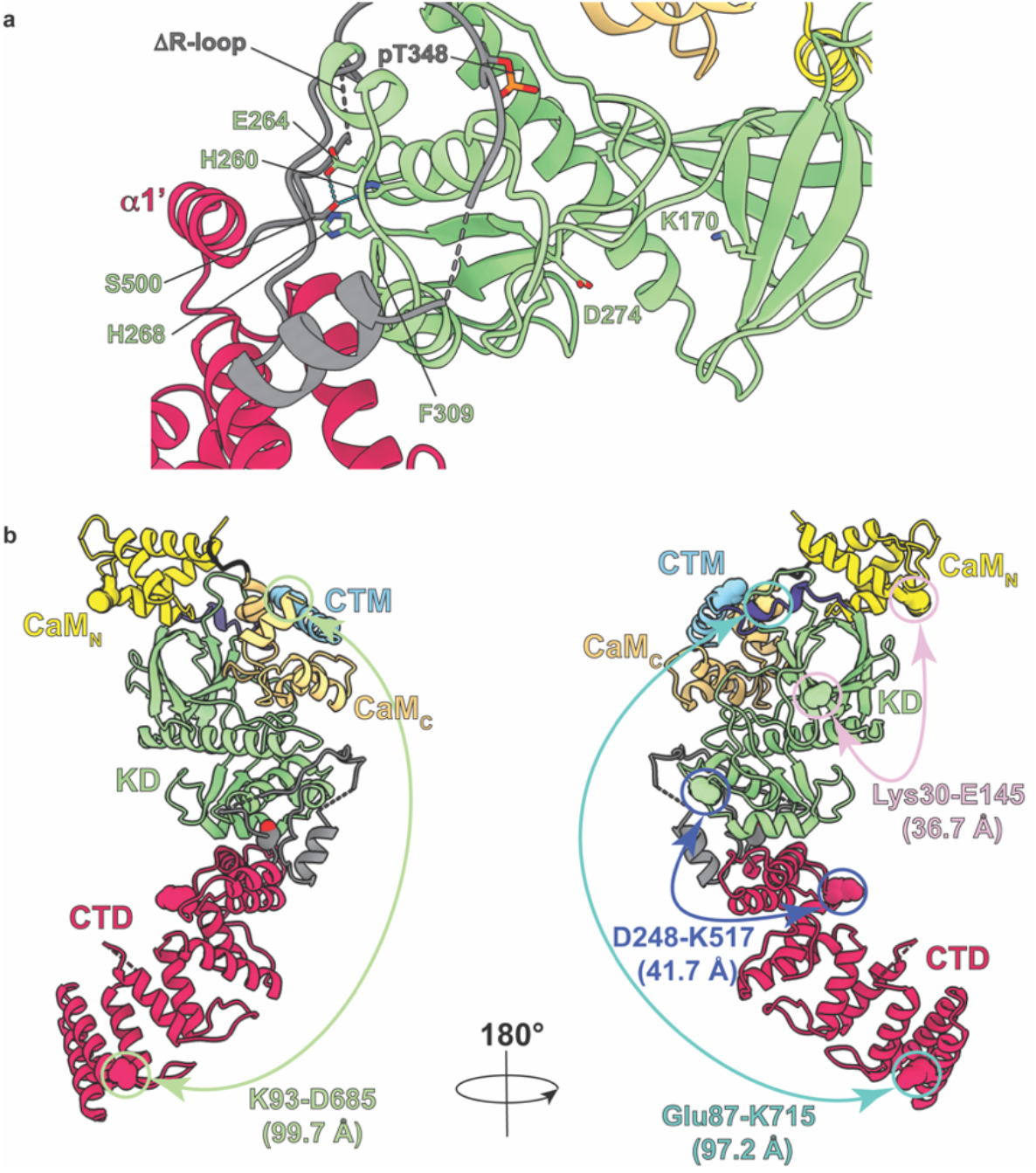
Auto-phosphorylation on S500 requires conformational changes. **a**, The secondary site of activating auto-phosphorylation (S500) is buried in a pocket within KD_C_, being stabilized through hydrogen bonds involving the side-chains of the Zn^2+^-coordinating H260, and E264, at a significant distance from the catalytic site (e.g., the S500,Oγ–D274,Oδ1 and the S500,Oγ–K170,Nζ distances are 20.0 and 29.7 Å, respectively; D274 is the putative catalytic base) suggesting that auto-phosphorylation at this position *in cis* would require significant conformational rearrangements including a partial unfolding of α1’. Also shown for reference is the position of *p*T348 bound at the PBP. **b**, Unambiguous intermolecular (between CaM and eEF-2K_TR_) and intramolecular (CTM/CTD, KD/CTD) crosslinks assigned for the CaM•eEF-2K_TR_ complex using the “zero-length” carbodiimide crosslinker EDC. While the two shorter Cα-Cα distances could be accommodated within a 1:1 complex through conformational flexibility, the longer distances would require substantial structural distortions to enable close contact of the KD with the CTD, and of the CTD with CaM, as proposed in an earlier computational model. However, all these crosslinks would be compatible with a 2:2 complex with an antiparallel assembly of the constituent CaM•eEF-2K_TR_ units. Such an arrangement would also facilitate *in trans* S500 auto-phosphorylation through smaller conformational changes. CaM and eEF-2K_TR_ residues are indicated by three- and one-letter codes, respectively.

**Extended Data Table S1.** Data collection and refinement statistics.

|  |  |
| --- | --- |
| **Data Collection** |  |
| Beamline | NSLS-II BEAMLINE 19-ID |
| Wavelength (Å) | 0.97946 |
| Space Group | P3<sub>1</sub>21 |
| Cell dimensions: |  |
| *a*, *b*, *c* (Å) | 58.50 58.50 365.78 |
| α, β, γ (°) | 90 90 120 |
| Resolution (Å) | 121.9 2.34 |
|  | (2.382, 2.342) |
| R<sub>merge</sub> | 0.362 (1.778) |
| *I*/σ(*I*) | 14.5 (2.5) |
| Completeness (%) | 100 (100) |
| Redundancy | 25.9 (24.5) |
| Wilson B-factor | 14.99 |
| Number of unique reflections | 32010 (1494) |
| **Refinement** |  |
| Resolution range (Å) | 50.66-2.34 |
| No. reflections | 31982 |
| R<sub>work</sub>/R<sub>free</sub> | 0.2293/0.2520 |
| Number of non-hydrogen atoms |  |
| Macromolecules | 5117 |
| Ligands | 8 |
| Solvent | 144 |
| Protein residues | 637 |
| RMS (bond) (Å) | 0.003 |
| RMS (angles) (°) | 0.533 |
| Ramachandran Plot |  |
| Most favored (%) | 91.1 |
| Additionally favored (%) | 8.7 |
| Generously favored (%) | 0.2 |
| Disallowed (%) | 0.0 |
| Rotamer outliers (%) | 0.38 |
| Clash-score | 4.29 |
| Average B-factor (Å<sup>2</sup>) | 35.37 |
| Macromolecules | 35.6 |
| Ligands | 29.35 |
| Solvent | 27.61 |
| PDB accession code | 7SHQ |

Values in parentheses are for the highest resolution shell. $R_{\text{merge}} = \sum |I - \langle I \rangle| / \sum I$ , where *I* is the observed intensity.

